# Exploring the genome and protein space of viruses

**DOI:** 10.1101/2022.11.05.515293

**Authors:** Congyu Lu, Yifan Wu, Zheng Zhang, Longfei Mao, Xingyi Ge, Aiping Wu, Fengzhu Sun, Yongqiang Jiang, Yousong Peng

## Abstract

Recent metagenomic studies have identified a vast number of viruses. However, the systematic assessment of the true genetic diversity of the whole virus community on our planet remains to be investigated. Here, we explored the genome and protein space of viruses by simulating the process of virus discovery in viral metagenomic studies. Among multiple functions, the power function was found to best fit the increasing trends of virus diversity and was therefore used to predict the genetic space of viruses. The estimate suggests that there are at least 8.23e+08 viral Operational Taxonomic Units (vOTUs) and 1.62e+09 viral protein clusters on Earth when assuming the saturation of the virus genetic space, taking into account the balance of costs and the identification of novel viruses. It’s noteworthy that less than 3% of the viral genetic diversity has been uncovered thus far, emphasizing the vastness of the unexplored viral landscape. To saturate the genetic space, a total of 3.08e+08 samples would be required. Analysis of viral genetic diversity by ecosystem yielded estimates consistent with those mentioned above. Furthermore, the estimate of the virus genetic space remained robust when accounting for the redundancy of sampling, sampling time, sequencing platform, and parameters used for protein clustering. This study provides a guide for future sequencing efforts in virus discovery and contributes to a better understanding of viral diversity in nature.

## Main text

Viruses are the most abundant and diverse biological entities on earth^1^. According to the Baltimore classification system, viruses can be classified into seven groups including double-stranded DNA viruses (dsDNA), single-stranded DNA viruses (ssDNA), double-stranded RNA viruses (dsRNA), positive-sense single-stranded RNA viruses (+ssRNA), negative-sense single-stranded RNA viruses (-ssRNA), positive-sense ssRNA viruses with a DNA intermediate(ssRNA-RT), and dsDNA viruses with ssRNA intermediates (dsDNA-RT), based on their genetic materials and replication mode^2^. Viruses exhibit wide variations in terms of host range, virion morphology, genome type, and size^3^. For example, the size of a virus genome can span from several hundred to millions of bases^4,5^. Beyond their impact on human health and diseases, viruses also play a crucial role in maintaining the balance of global ecosystems. Previous studies have suggested that viruses are responsible for the daily demise of 20% of bacterial hosts in the oceans^1^.

An enormous number of viruses have been identified, thanks to the rapid development of Next-Generation-Sequencing (NGS) technology, in comparison to traditional virus identification methods based on virus isolation^6-8^. For example, the IMG/VR database, recognized as the largest virus sequence database, has amassed over 15 million viral genomic sequences^6^. Furthermore, novel viruses continue to be discovered at an unprecedented rate. The Global Ocean Viromes (GOV) 2.0 project, for instance, has unveiled nearly 200,000 marine viral populations through the sequencing of 145 ocean samples^7^. This raises a natural question regarding the genetic space of viruses. Previous studies by Forest Rohwer estimated that there were 100 million phage species on Earth^9^. The Global Virome Project put the estimate at 1.67 million viruses in birds and mammals^10^. Both studies based their estimates on the assumption that a host species harbors, on average, dozens of viruses, without considering the actual process of virus discovery through sampling and sequencing. This study delves into the genome and protein spaces of viruses by simulating the process of virus discovery in viral metagenomic studies.

### The increasing trends of the virus genome and protein space

Firstly, we investigated the increasing trends in the virus genome and protein space as the number of samples increased. Specifically, we randomly selected 10 sequenced samples from the IMG/VR database without replacement, and then counted the accumulated number of viral Operational Taxonomic Units (vOTUs) and viral Protein Clusters (vPCs) (the definitions of vOTU and vPC are described in the Methods section) identified in the accumulated samples selected up to that point. This process was repeated until no samples remained (see Methods). The entire selection process was repeated 100 times. As depicted in Figure 1A, both the number of vOTUs and vPCs increased rapidly. With all the samples in the IMG/VR database, a total of 7,721,789 vOTUs and 40,464,268 vPCs were identified.

**Figure 1.**
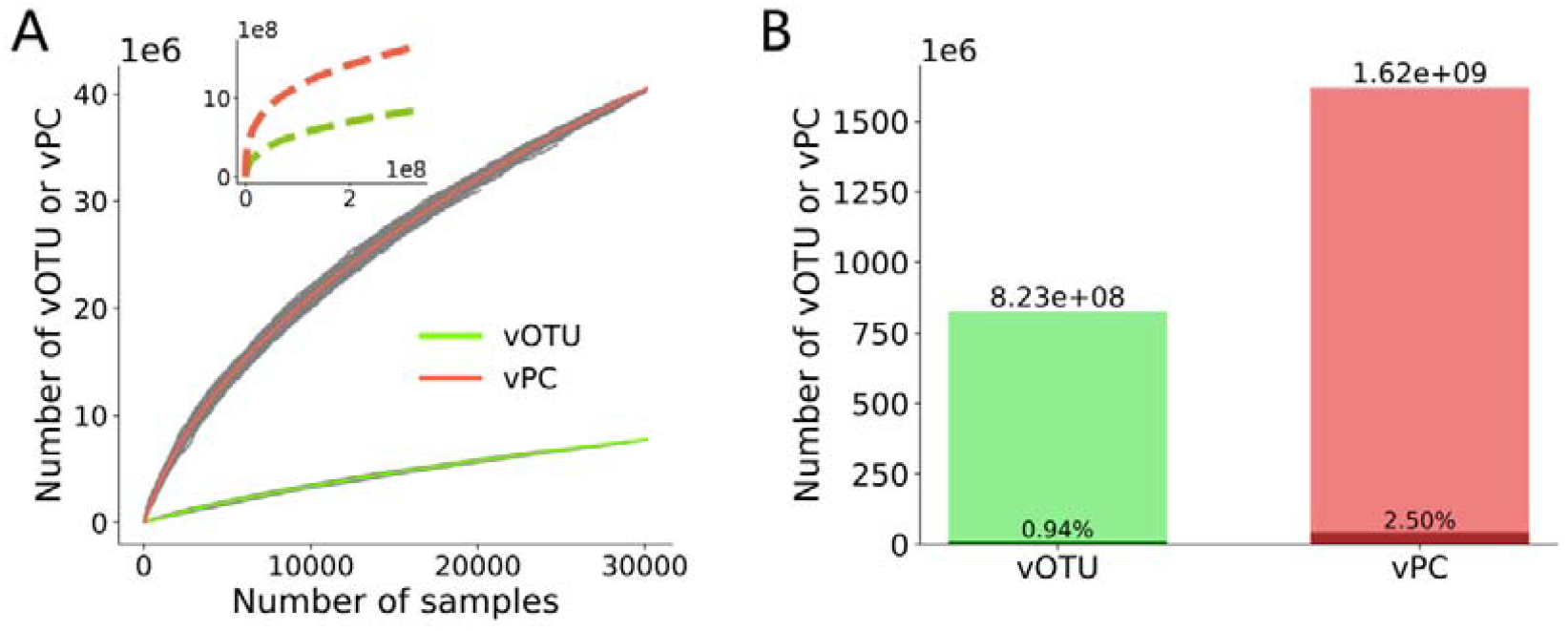
Estimating the virus genome and protein space. (A) Fitting the increasing trend of vOTUs and vPCs versus the number of selected samples. The gray points represent the number of vOTUs and vPCs identified in the selected samples in 100 simulations of sequencing studies (see Methods). The solid line depicts the fitted curve of the Power function for vOTUs (in green) and vPCs (in red). The top-left sub-figure illustrates the prediction of the Power function. (B) The estimated total number of vOTUs and vPCs when the virus genetic space was saturated, along with the proportion of explored viral genetic space at levels of vOTUs and vPCs.

To capture the increasing trends in the viral genetic space, we employed several common mathematical functions, including Power, Sigmoid, Second-order polynomial, Triple-order polynomial, Logarithmic, Exponential, and Inverse proportional, to fit the relationship between the number of vOTUs or vPCs and the number of samples selected. To determine the best function, 3000 samples were randomly selected as the training data, while the remaining 27,158 samples were used to test these functions. As illustrated in Figure S1, only the Power function fitted well for both vOTU and vPC, achieving a much lower Mean Squared Error (MSE) than other functions on the testing samples, even though all functions fit well on the training samples. Similar results were obtained when changing the dataset size for training and testing, with the Power function consistently fitting best among all functions (see Table S1). Therefore, the Power function was used to predict the increasing trends of vOTU and vPC, assuming that more sequenced samples would be added in future studies (refer to the top-left sub-figure in Figure 1A). According to the predictions, vOTUs and vPCs would continue to increase rapidly and then begin to expand with a slowing rate.

While the generation and disappearance of viruses occur daily due to their rapid evolution, there exists a balance between the birth and death of viruses, suggesting that the number of viruses on Earth should be finite. Consequently, it is anticipated that the identification of novel viruses would decrease as sampling continues. However, given the vast diversity and rapid evolution of viruses, it may be challenging to identify all viruses on Earth. To estimate the total number of viruses, the virus genetic space was assumed to be saturated when less than one novel virus was identified in a sample. Based on this assumption, the total numbers of vOTUs and vPCs were estimated to be 8.23e+08 and 1.62e+09, respectively, when the virus genetic space was saturated (see Figure 1B). The Proportion of Explored Viral Genetic Space (PEVGS), defined as the total number of vOTUs or vPCs identified in the IMG/VR database divided by the estimated numbers when the virus genetic space was saturated, was calculated to be 0.94% for vOTU and 2.50% for vPC. As depicted in Figure 1A, the number of new viruses identified in new samples decreases as sampling continues. It was estimated that 3.08e+08 samples were required to saturate the genetic space. Unfortunately, only less than 40,000 samples had been sequenced so far according to the IMG/VR database.

### Taxonomy composition of viruses in different ecosystems

According to the GOLD (Genomes OnLine Database) ecological classification system^11^, the samples used in this study primarily originated from seven ecosystems, including two environmental ecosystems (Aquatic and Terrestrial ecosystems accounting for 44% and 23% of all samples, respectively), three host-associated ecosystems (Mammal, Other-animal, and Plant ecosystems accounting for 13%, 2%, and 5% of all samples, respectively), and two engineered ecosystems (Wastewater and Built-environment ecosystems accounting for 1% and 4% of all samples, respectively) (refer to Table S2). The investigation into the taxonomic composition of viruses in different ecosystems involved calculating the proportion of vOTUs of different classes in each ecosystem^12^. As depicted in Figure 2A, the taxonomic composition of viruses across different ecosystems exhibited similarity, with the class of *Caudoviricetes* and unclassified viruses collectively constituting approximately 90% of all vOTUs. The class of *Caudoviricetes* held the largest proportion among all viral classes in every ecosystem, ranging from 71% to 96%. The taxonomic composition of the remaining vOTUs (approximately 10%) varied significantly among ecosystems. For instance, the class of *Faserviricetes* dominated the remaining vOTUs in the Built-environment ecosystem, where few other viruses were detected. Conversely, in the Other-animal ecosystem, the class of *Polintoviricetes* prevailed among the remaining vOTUs. Interestingly, the Aquatic ecosystem exhibited an enrichment of the class *Megaviricetes*, aligning with the predominance of their hosts (including algae and amoeba) in this ecosystem^12^

**Figure 2.**
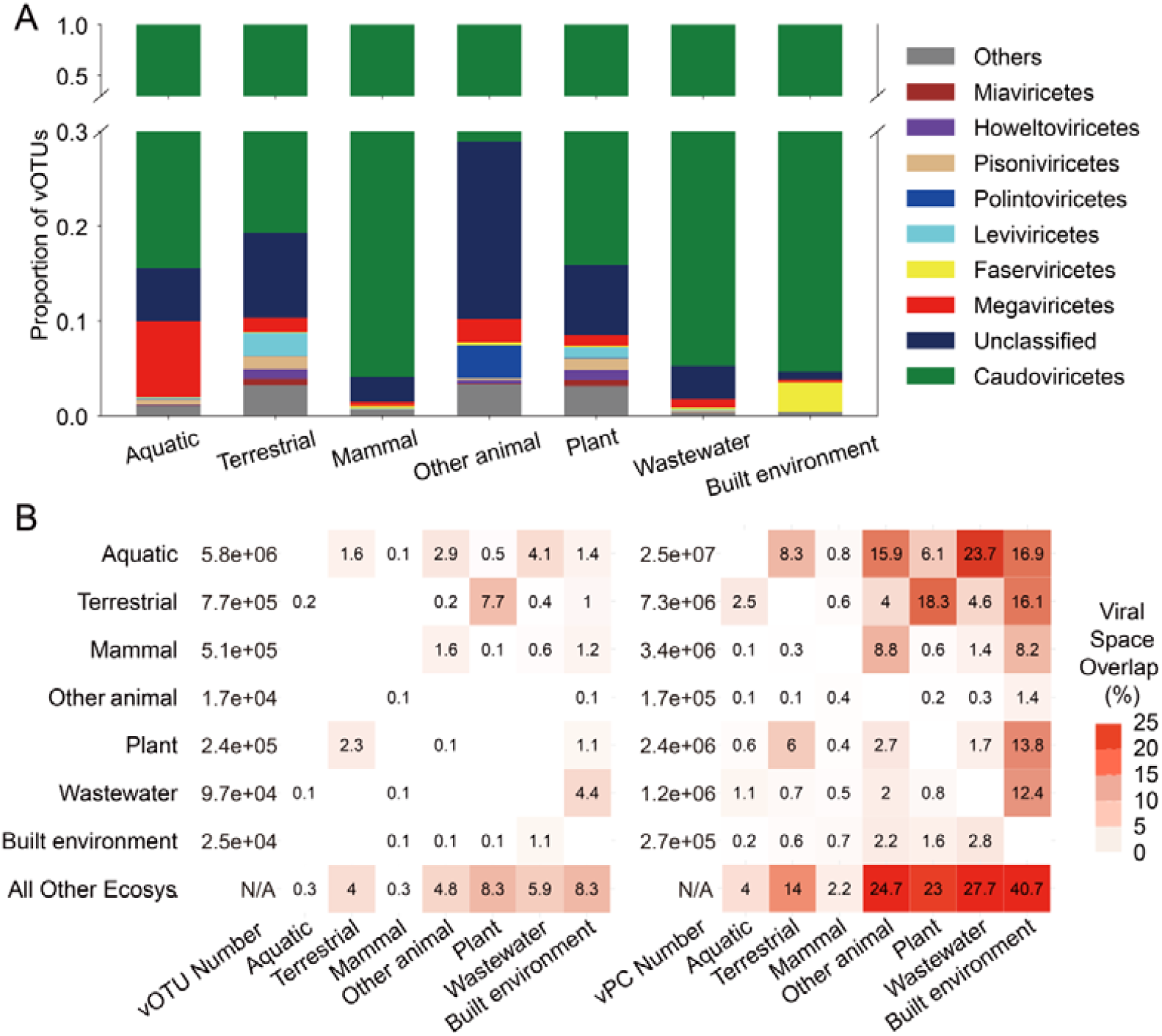
Analysis of the viral genome and protein space by ecosystem. (A) The taxonomic composition of viruses identified in different ecosystems. (B) The proportion of vOTUs and vPCs shared between different ecosystems. The numbers presented in the heatmap indicate the percentage of vOTUs or vPCs shared between two ecosystems (indicated on the left and bottom) relative to the total vOTUs or vPCs in the ecosystem shown at the bottom.

### Shared genetic diversity among different ecosystems

We then proceeded to analyze the extent of shared viral genetic diversity among different ecosystems. As illustrated in Figure 2B, various ecosystems exhibited a very small proportion of shared vOTUs. The proportion of shared vOTUs between any given ecosystem and other ecosystems ranged from 0.3% to 8.3%. Notably, the Aquatic ecosystem, although possessing the largest number of vOTUs, only covered a very small proportion of vOTUs in other ecosystems. For example, it covered 0.1% and 0.5% of vOTUs in Mammal and Plant ecosystems, respectively. Interestingly, the Plant ecosystem exhibited the highest degree of sharing with the Terrestrial ecosystem.

In contrast, the proportions of shared vPCs between different ecosystems were considerably larger than those of the shared vOTUs, ranging from 2.2% to 40.7%. The Mammal ecosystem had the smallest proportion of shared vPCs with other ecosystems (2.2%). Moreover, more than one-fifth of vPCs in three ecosystems, namely Other-animal, Plant, and Wastewater, were shared with other ecosystems. Remarkably, for the Built-environment ecosystem, over 40% of the vPCs were shared with other ecosystems.

### The virus genome and protein space estimated by ecosystem

To analyze the increasing trend of virus genetic diversity by ecosystem as the number of samples increases, random sampling was conducted for each ecosystem, utilizing only samples from the corresponding ecosystem (refer to Table S2 for the number of samples in different ecosystems). The increasing trends of vOTUs and vPCs in each ecosystem were analyzed and fitted using Power functions. As depicted in Figure 3A, the increasing trends of vOTUs in all ecosystems exhibited similarity, showing a slowing rate as the number of selected samples increased. Notably, the number of vOTUs in the Aquatic ecosystem increased most rapidly with the growing number of selected samples, while the increase in other ecosystems was considerably slower. At the vPC level (Figure 3B), the overall increasing trends mirrored those observed at the vOTU level. However, the differences in increasing trends of vPCs between the Aquatic and other ecosystems, especially the Aquatic and Terrestrial ecosystems, were smaller than those observed at the vOTU level.

**Figure 3.**
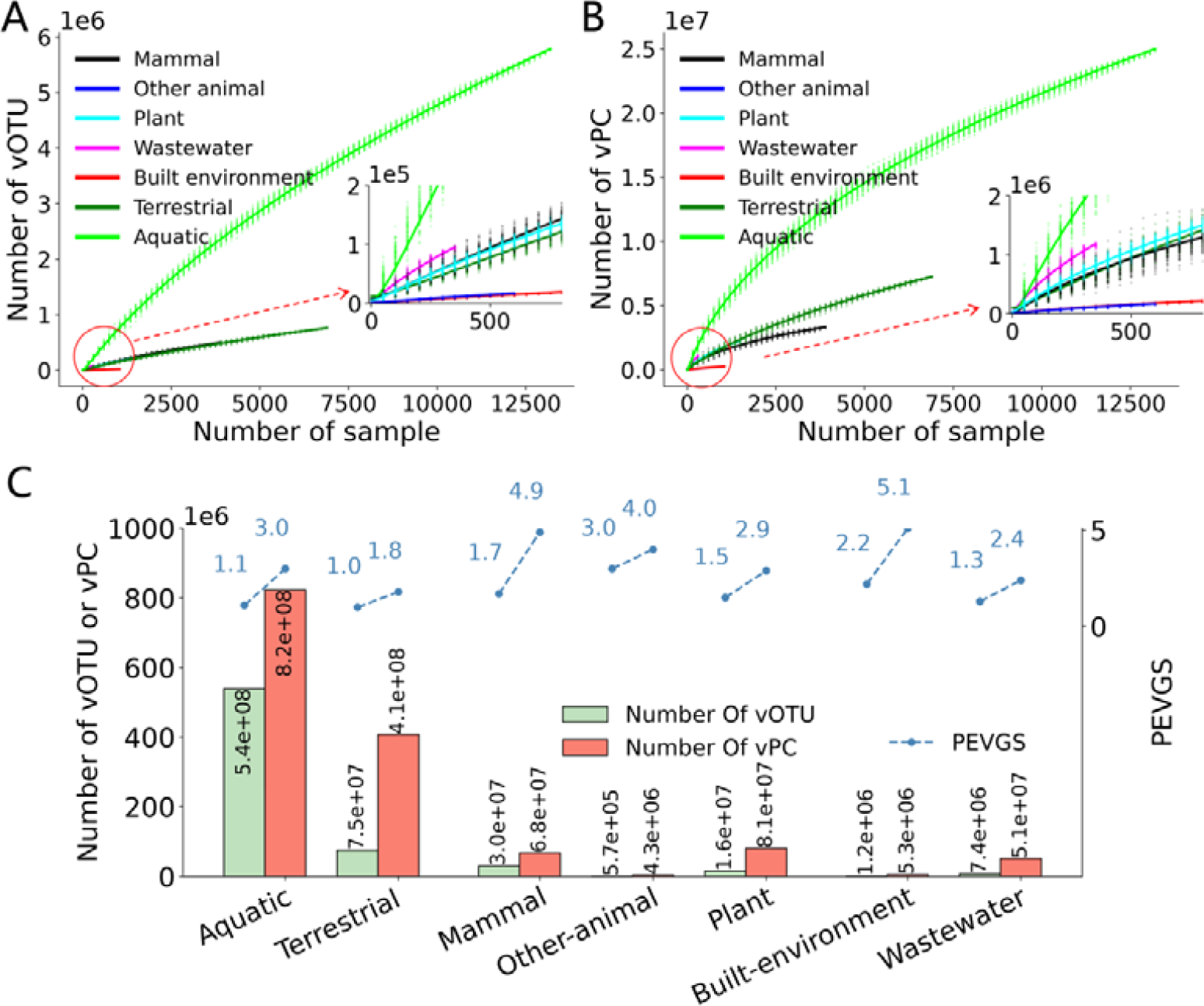
Analysis of the Viral Genome and Protein Space by Ecosystem. (A) and (B) refer to the increasing trends of the numbers of vOTUs and vPCs, respectively, as the number of selected samples in different ecosystems. The simulation process was conducted based on the ecosystem. (C) The histogram displays the estimated total numbers of vOTUs and vPCs in each ecosystem when the virus genetic space was saturated. Additionally, the Proportion of Explored Viral Genetic Space (PEVGS) of viruses in each ecosystem is illustrated at levels of vOTU and vPC (indicated by blue dashed lines).

We then estimated the total numbers of vOTUs and vPCs in different ecosystems when the viral genetic space was saturated (refer to Figure 3C). Predictions indicated that the Aquatic ecosystem was expected to have the largest number of vOTUs (5.4e+08) and vPCs (8.2e+08), followed by the Terrestrial ecosystem (7.5e+07 vOTUs and 4.1e+08 vPCs) and the Mammal ecosystem (3.0e+07 vOTUs and 6.8e+07 vPCs). In contrast, the four remaining ecosystems were predicted to contain considerably less viral diversity, with the Built-environment and Wastewater ecosystems projected to have only 1.2e+06 and 7.4e+06 vOTUs, respectively.

The PEVGS was calculated for each ecosystem at levels of vOTU and vPC (depicted in Figure 3C). Across all ecosystems, PEVGS values increased from the level of vOTU (ranging from 1.0% to 3.0%) to vPC (ranging from 1.8% to 5.1%). Notably, host-associated ecosystems exhibited higher PEVGS values compared to environmental and engineered ecosystems. For instance, the Mammal ecosystem had PEVGS values of 1.7% and 4.9% at levels of vOTU and vPC, respectively, while those for the Terrestrial ecosystem were 1.0% and 1.8%, respectively.

### Stability analysis of the estimate of the virus genetic space

Finally, we delved into the influence of various factors on the estimate of the virus genetic space. The first factor considered was the redundancy of samples, where one sample might undergo sequencing multiple times or the sequencing data of the sample might be utilized in multiple studies. Upon eliminating redundancy using the NCBI BioSample, 25,836 biosamples were retained, and the estimated number of virus vOTUs was 9.44e+08. Similarly, when redundancy was removed using the NCBI BioProject (ensuring only one sample was used in each BioProject), the estimated number of virus vOTUs was 9.40e+08. Notably, both estimates after eliminating redundancy closely aligned with the original estimate (8.23e+08) (refer to Table 1), suggesting that sample redundancy had a minor influence on the estimate of the virus genetic space.

**Table 1.** Influence of different factors on the estimate of virus genetic space.

| Part I: Redundancy of samples |  |  |
| --- | --- | --- |
|  | Number of samples used | Estimated number of vOTUs |
| Unique IMG sample | 30,158 | 8.23e+08 |
| Unique BioSample | 25,836 | 9.44e+08 |
| Unique BioProject | 20,260 | 9.40e+08 |
| Part II: Sampling time |  |  |
| Before 2018 | 15,349 | 6.15e+08 |
| 2018 and after | 14,809 | 8.38e+08 |
| Part III: Sequencing platform |  |  |
| Illumina NovaSeq | 7,426 | 5.38e+08 |
| Illumina HiSeq | 15,546 | 5.60e+08 |

The number of viruses identified in samples varied significantly over different periods, experiencing a substantial increase in 2018 (see Figure S2). To investigate the impact of the year of virus identification on the estimate, all samples used in the study were categorized into two groups: one group comprised samples obtained before 2018, totaling 15,349 samples, while the other group included samples obtained in 2018 and later, totaling 14,809 samples. The virus genetic space was then estimated for each group of samples. As outlined in Table 1, the estimated numbers of viral vOTUs for the two groups were 6.15e+08 and 8.38e+08, respectively. Despite the former showing a nearly 30% decrease compared to the original estimate, both estimates were of the same order of magnitude as the original estimate.

The number of viruses identified from sequencing data generated by different sequencing platforms exhibited considerable variation (see Figure S3). For instance, a median of 140 and 93 vOTUs were identified per sample sequenced by Illumina NovaSeq and HiSeq, respectively, while only 17 vOTUs were identified per sample sequenced by Illumina MiSeq. Consequently, we investigated the influence of the sequencing platform on the estimate of virus genetic space. As the majority of samples were sequenced by Illumina NovaSeq (7,426, 25%) and HiSeq (15,546, 52%), we estimated the virus genetic space based on samples sequenced exclusively by Illumina NovaSeq or HiSeq, resulting in estimated values of 5.38e+08 and 5.60e+08 vOTUs, respectively (refer to Table 1). Once again, both estimates were of the same order of magnitude as the original estimate.

Virus protein clusters were obtained by clustering protein sequences with the threshold of identity and coverage set at 0.7. We then investigated the influence of these thresholds on the estimate of viral genetic space. Increasing the threshold from 0.5 to 0.8 resulted in an increase in the number of vPCs from 3.24e+07 to 4.66e+07. The estimated total number of vPCs also increased from 1.10e+09 to 2.01e+09, remaining within the same order of magnitude as the original estimate when the threshold was set to 0.7 (refer to Table 2).

**Table 2.** The influence of thresholds of identity and coverage on the estimate of vPCs.

| Threshold for identity and coverage | Number of vPCs obtained | Estimate of the number of vPCs |
| --- | --- | --- |
| 0.5 | 3.24e+07 | 1.10e+09 |
| 0.6 | 3.59e+07 | 1.30e+09 |
| 0.7 | 4.05e+07 | 1.62e+09 |
| 0.8 | 4.66e+07 | 2.01e+09 |

## Discussion

In recent years, numerous large-scale projects targeting the entire virome in various specific ecological environments have uncovered an abundance of novel viruses. Despite these efforts, a comprehensive understanding of the ultimate extent of virus genetic diversity on Earth has remained elusive. This study represents the first attempt to estimate the potential virus genome and protein space based on current virus genetic diversity, revealing a total of 8.23e+08 vOTUs and 1.62e+09 vPCs on Earth. Notably, the estimated virus genetic space aligned with the sum of viral genetic diversity estimated individually within each ecosystem. Furthermore, the estimates of virus genetic space remained relatively stable when considering multiple factors, highlighting the robustness of the estimations regarding viral genome and protein space.

However, it’s crucial to note that less than 3% of the viral genetic space has been unveiled thus far, emphasizing the necessity for additional virome projects to comprehensively capture viral genetic diversity. Unfortunately, as more viruses are identified, the number of novel viruses in new samples diminishes. This study estimates that a substantial 3.08e+08 samples would be required to encompass the entire viral genetic diversity. Notably, samples from the Aquatic ecosystem emerged as key contributors, capturing a significantly larger portion of viral genetic diversity than other ecosystems and should thus be prioritized in virus discovery efforts. Moreover, certain viral orders exhibited preferences for specific ecosystems, exemplified by the enrichment of Megaviricetes in the Aquatic ecosystem. Sequencing studies tailored to such preferences hold promise for the precise identification of specific kinds of viruses.

Despite utilizing the ICTV and Baltimore systems for viral taxonomy, most viruses analyzed in this study remained unclassified at the order and family levels. A critical need exists to develop an expandable classification framework for all viruses. The ideal classification framework should have the ability to classify all viruses, easy to use, and scalable. Previous studies by Jang and colleagues have developed the vConTACT for classification of prokaryotic viruses based on genomic sequences^13^. In the study, a gene sharing network of viruses based on the shared protein clusters among viral genomes was built, and a distance-based clustering and metric were integrated to provide the measures of clustering confidence. vConTACT is a scalable and automated classification tool as it can build classifications for more than 10,000 sequences which were previously unclassified in oceanic viromics studies. Moreover, Jang’s study provided a novel and important framework for organizing viral genetic diversity, and demonstrated that the gene content-based methods are suitable for a unified classification of all viruses. Unfortunately, it was not widely used in virome studies. There was still a lack of a feasible classification framework for the whole virome on Earth.

Several limitations must be acknowledged. Firstly, the estimated virus protein space may be influenced by parameter settings in protein clustering. Fortunately, the estimated total number of vPCs exhibited stability across varying parameters, ensuring the robustness of the estimate. Secondly, the study predominantly sampled from seven ecosystems, with potential bias in their representation. Some ecosystems, such as Other-animal and extreme environments, might be underrepresented, impacting the estimate of viral genetic diversity. Additionally, regional and country-based sampling bias, favoring developed countries, may further affect the accuracy of the estimate. Lastly, the RNA viruses were under-represented as approximately 80% of the samples collected in the IMG/VR database were sequenced using the metagenomic strategy that favors DNA viruses. In combination, it’s essential to view the estimate of virus genetic space as a conservative lower bound.

## Conclusion

In summary, this study explores the virus genome and protein space, estimating the total number of vOTUs and vPCs on Earth when the virus genetic space is saturated. It serves as a guiding framework for future virus discovery sequencing efforts and significantly contributes to our understanding of viral diversity in nature.

## Methods Data retrieval

The viral genome and protein sequences were retrieved from the IMG/VR database (v4)^6^. Meta information for metagenomic samples, from which viruses were identified, was obtained from the IMG/M database^14^. Samples derived from experimental cultures were excluded as they do not adequately reflect natural virus diversity. Restricted samples were removed according to the JGI data utilization policy^6^. A total of 30,158 samples were retained for further analysis, encompassing 13,746,160 viral genomic sequences, 7,721,789 vOTUs (defined as clusters of genomic sequences with an average of 95% or higher sequence identity on more than 85% of genomic regions according to the IMG/VR database^6^), and 179,131,383 protein sequences.

### Generation of protein clusters

To reduce computational costs in generating protein clusters, all viral protein sequences were initially clustered using the linclust algorithm of MMseqs2, with both coverage and identity set to 0.7^15^. This process yielded a total of 40,464,268 vPCs. The influence of parameter selection on the generation of vPCs is detailed in Table 2.

### Simulation of sampling in viral metagenomic studies

To simulate the virus discovery process in viral metagenomic studies, 10 sequenced samples were randomly selected from all samples without replacement. The number of vOTUs and vPCs identified in the accumulated samples selected up to that point was recorded. This sampling process was iterated until no samples remained. The increasing trend of the accumulated vOTUs and vPCs versus the number of accumulated samples selected was analyzed. The simulation process was repeated 100 times to mitigate randomness in sampling.

### Predicting the size of vOTUs and vPCs when the genetic space was saturated

Various mathematical functions, including Sigmoid, Power, Exponential, Second-order Polynomial, Triple-order polynomial, Logarithmic, and Inverse Proportional functions, were employed to fit the increasing trend of the accumulated vOTUs versus the common logarithm of the accumulated samples selected in the simulation process. This was achieved using the scipy.optimize.curve_fit package in Python (formulas for these functions listed in Supplementary Materials)^18^.

## Ethics approval and consent to participate

Not applicable

## Consent for publication

All authors gave their consent for publication.

## Availability of data and materials

All data used in the study are derived from the IMG/VR and IMG/M databases.

## Authors’ contributions

Y.P. conceived and designed the analysis. Y.P., C.L. and Y.W. performed the analysis and prepared all figures. Y.P., C.L., Z.Z., Y.J., F.S., L.D., A.W., X.G. and L.M. wrote the main manuscript text. All authors reviewed the manuscript.

## Acknowledgements

We would like to thank Prof. Simon Roux for guidance of the usage of IMG/VR database, and thank members in Peng’s lab for helpful suggestions.

