## Supplementary material for "Exploring the genome and protein space of viruses": Figure S

**Mathematical functions used in the study**

Several mathematical functions including Sigmoid (denoted as S(x)), Power (denoted as Power(x)), Second-order polynomial (denoted as Ploy_2_(x)), Triple-order polynomial (denoted as Ploy_3_(x)), Logarithmic (denoted as L(x)), Exponential (denoted as E(x)), and Inverse proportional (denoted as I(x)) functions were used to fit the increasing trend of the number of accumulated vOTUs versus the common logarithm of the accumulated number of samples selected (defined as x). The formulas for these functions were listed as follows. The “a”, “b”, “c” and “d” were parameters of functions.

$S\left( x \right)=\frac{c}{1+e^{-ax+b}}$	（1）

$Power\left( x \right)={ax}^{b}+c$	（2）

${Ploy}_{2}\left( x \right)={ax}^{2}+bx+c$	（3）

${Ploy}_{3}\left( x \right)={ax}^{3}+{bx}^{2}+cx+d$	（4）

$L\left( x \right)=a{log}_{10}^{x}+b$	（5）

$E\left( x \right)=ae^{-bx}+c$	（6）

$I\left( x \right)=\frac{a}{x}+b$	（7）

**Supplementary Figures**

**Figure S1**. Fitting the increasing trends of vOTUs (A) and vPCs (B) with different mathematical functions. 3000 samples were used in training (red points), while the remaining 27,158 samples were used in testing (blue points). The MSEs of each function on the testing samples were listed for each function on the legend.

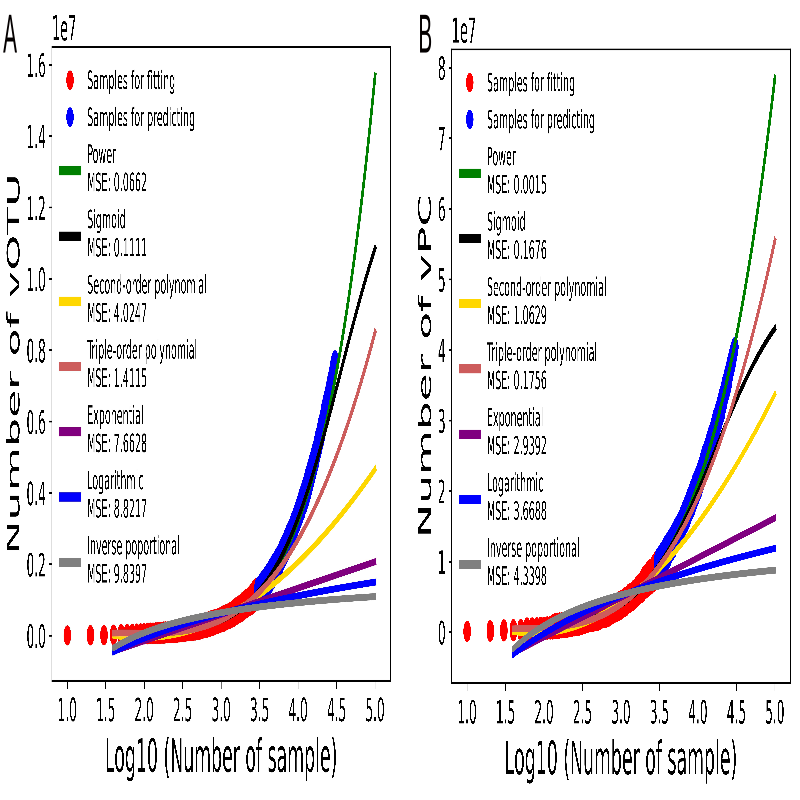

**Figure S2.** Number of vOTUs identified per sample for samples obtained in years from 2010 to 2022.

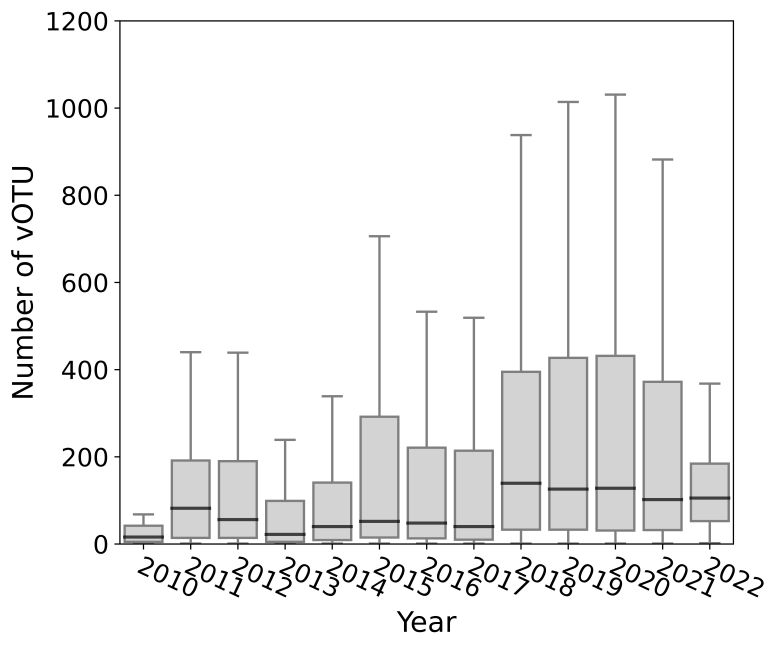

**Figure S3**. The number of vOTUs identified per sample for data generated by different sequencing platforms.

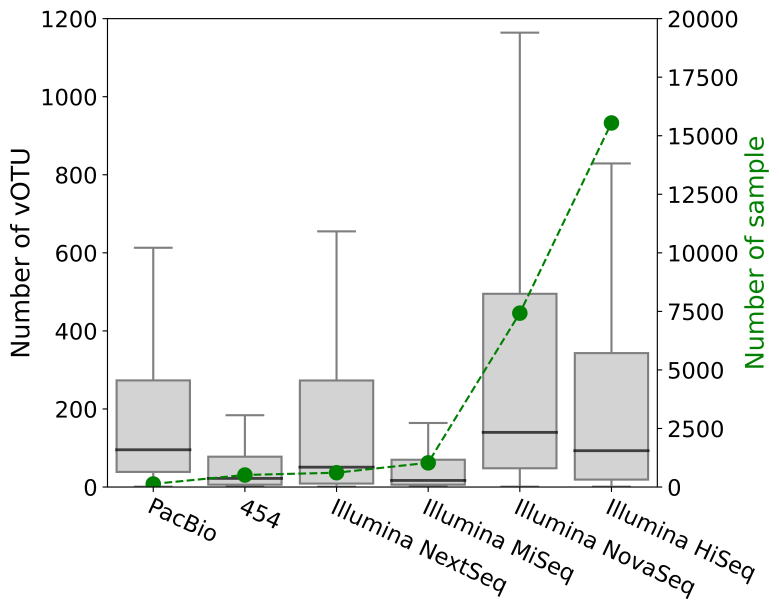

**Supplementary Tables**

**Table S1**. The Mean Squared Error (MSE) of mathematical functions on the testing samples when changing the size of training and testing samples. The smallest MSEs were highlighted in bold for each set of training samples.

|  | **Number of training samples** | **Power** | **Sigmoid** | **Second-order polynomial** | **Triple-order polynomial** | **Exponential** | **Logarithmic** | **Inverse poportional** |
| --- | --- | --- | --- | --- | --- | --- | --- | --- |
| vOTU | 500 | **4.8E+01** | 8.2E+01 | 2.0E+02 | 1.2E+02 | 2.6E+02 | 2.7E+02 | 2.8E+02 |
|  | 1000 | **5.4E+00** | 8.8E+01 | 4.6E+01 | 2.4E+01 | 6.8E+01 | 7.3E+01 | 7.7E+01 |
|  | 2000 | **4.9E-01** | 2.7E+01 | 1.0E+01 | 4.2E+00 | 1.7E+01 | 1.9E+01 | 2.1E+01 |
|  | 3000 | **6.6E-02** | 1.1E-01 | 4.0E+00 | 1.4E+00 | 7.7E+00 | 8.8E+00 | 9.8E+00 |
|  | 5000 | **8.5E-04** | 4.2E-03 | 1.1E+00 | 2.9E-01 | 2.6E+00 | 3.2E+00 | 3.7E+00 |
|  | 7000 | **9.1E-04** | 8.8E-03 | 4.6E-01 | 9.4E-02 | 1.2E+00 | 1.6E+00 | 1.9E+00 |
| vPC | 500 | **4.0E+00** | 9.5E+01 | 4.3E+01 | 1.6E+01 | 7.1E+01 | 7.8E+01 | 8.2E+01 |
|  | 1000 | **6.4E-01** | 8.7E+00 | 1.2E+01 | 3.9E+00 | 2.3E+01 | 2.7E+01 | 2.9E+01 |
|  | 2000 | **1.8E-02** | 5.9E-01 | 2.7E+00 | 6.0E-01 | 6.5E+00 | 7.7E+00 | 8.8E+00 |
|  | 3000 | **1.5E-03** | 1.7E-01 | 1.1E+00 | 1.8E-01 | 2.9E+00 | 3.7E+00 | 4.3E+00 |
|  | 5000 | **3.7E-03** | 2.2E-02 | 3.1E-01 | 3.2E-02 | 1.1E+00 | 1.5E+00 | 1.8E+00 |
|  | 7000 | **1.9E-03** | 1.2E-02 | 1.3E-01 | 1.0E-02 | 5.5E-01 | 7.6E-01 | 1.0E+00 |

**Table S2.** The number of samples in different ecosystems.

| **Ecosystem Type** | **Ecosystem** | **Number of sample**s |
| --- | --- | --- |
| Environmental | Aquatic | 13,230 |
| Environmental | Terrestrial | 6,998 |
| Environmental | Others | 64 |
| Host-associated | Mammal | 4,016 |
| Host-associated | Other-animal | 613 |
| Host-associated | Plant | 1,630 |
| Host-associated | Others | 424 |
| Engineered | Wastewater | 361 |
| Engineered | Built-environment | 1,103 |
| Engineered | Others | 1,719 |
| Total | | 30,158 |
